# Neotropical birds and mammals show divergent behaviour responses to human pressure

**DOI:** 10.1101/2022.09.22.509119

**Authors:** Pablo Jose Negret, Mathew Scott Luskin, Bibiana Gomez-Valencia, Angelica Diaz-Pulido, Luis Hernando Romero, Adriana Restrepo, Julie G. Zaehringer, Jose Manuel Ochoa-Quintero, Calebe Pereira Mendes

## Abstract

Human presence and habitat disturbance (together ‘human pressure’ hereafter) can generate a deep fear in animals and this can influence their behaviour. Altered animal behaviour, such as shifts in diel activity patterns, affect many species and species interactions, which can induce changes in individual fitness, species-level population persistence, evolutionary dynamics, and ecosystem-level biodiversity. However, whether dial activity behavioural responses to human pressure are consistent among key functional groups has been poorly studied. For example, while medium to large mammal species tend to become more nocturnal in areas with high human pressure, it’s unclear if sympatric/co-occurring birds display similar or opposite patterns. This is an important knowledge gap because synchronous or opposing guild-level shifts can shape consequences for food-web dynamics (predation and competition), stability of interaction networks and ecosystem functioning. Here we used information from camera trapping along a gradient of human pressure in the Colombian Llanos region to assess diel activity changes in birds and mammals. We found that the diel activity of over 45% of the bird and 50% of the mammals assessed significantly changed where there was higher human pressures, with mammals becoming more nocturnal and birds more diurnal. The opposing behavioural responses to humans among vertebrate functional groups has significant repercussions for the fields of community ecology, including intraguild predation and competition, and may be a significant ecosystem-level conservation consideration.

## Introduction

The extent of human pressure has increased dramatically in the last 20 years with over two thirds of the global land area under moderate or intense human pressure (Williams et al., 2020b). The related habitat changes and specific anthropic pressures (e.g. hunting) have vastly altered wildlife communities including extirpations, extinctions, and changing ecosystem functioning (Dirzo et al., 2014; Haddad et al., 2015). These direct effects are also accompanied by the indirect effects of humans on the behaviour of remaining wildlife, which poses a significant threat to species fitness and species interactions as well as overall biodiversity and ecosystem functioning (Gaynor et al., 2018; Sih et al., 2011; Zanette and Clinchy, 2019).

Human presence and habitat disturbance (together ‘human pressure’ hereafter) can generate a deep fear in animals and this “landscape of fear” can influence their diel activity patterns and use of space (Gallagher et al., 2017; Laundre et al., 2010; Zanette and Clinchy, 2019). Medium and large mammals tend to avoid humans by increasing their nocturnality (Gaynor et al., 2018; Mendes et al., 2020) and reducing their home range closer to urban environments (O’Donnell and Delbarco-Trillo, 2020). However, a recent study found scarce evidence of activity pattern shifts for mammals in areas with high human pressure (oil palm plantations) in the Colombian Llanos (Pardo et al., 2021). Further investigation is needed since these variations in wildlife responses to humans are not well understood but they could have immense repercussions for species interactions such as predation and competition, which can restructure entire food webs, energy flows and ecosystem functioning.

Bird behavioural responses to humans may differ from mammals but our understanding is more limited, especially in the tropics (Bonier et al., 2007; Chace and Walsh, 2006; Zanette and Clinchy, 2019). Chilean forest birds shifted their daily activity peaks from early morning in old-growth forests, to noon in plantations, and to the afternoon in logged forests (Fontúrbel et al., 2021) and forest birds in Hawaii showed increased diurnality in fragmented landscapes (Smetzer et al., 2022). One difference underlying variation in vertebrate functional group responses could be related to harvest rates, which are often predicted by larger body size and ease of capture (less arboreality or flight) (Dirzo et al., 2014). In a global review of hunting in tropical forests, birds and mammals both showed significant declines in hunting-accessible areas, although 25% less for birds than mammals (Benítez-López et al., 2017). There is also evidence that birds need to forage more in fragmented habitats, which could increase their diurnality (Redpath, 1995; Saunders, 1980). Fearful animals near humans may perceive safety differently depending on their ability to hide versus escape, with arboreal birds preferring higher daytime visibility and ease of escape via flight, while terrestrial mammals may view nocturnal stealthy foraging as safer (Gallagher et al., 2017; Zanette and Clinchy, 2019). Understanding bird behavioural responses to human pressure and how this responses may differ from those of mammals is an important knowledge gap because synchronous or opposing guild-level shifts among mammals and birds can shape consequences for food-web dynamics and ecosystem functioning (Zanette and Clinchy, 2019).

Colombia, one of the most biodiverse countries is affected by high levels of habitat degradation and human pressures have doubled in the last 50 years (Correa Ayram et al., 2020). However, there is limited evidence if there are changes to wildlife behaviour do to this increase in human pressure. We used camera trapping along a gradient of human pressure in the Colombian Llanos to measure changes in daily activity patterns of birds and mammals. For medium and large birds and mammals, we expected to observe shifts towards nocturnality for species susceptible to poaching and hunting and no changes for small species. Finally, we expect to see congruent responses from birds and mammals with similar ecological characteristics, such as game species, and species with similar body size and diet.

## Methods

### Survey

To detect possible shifts in the activity patterns of birds and mammals, we used a large dataset of camera-traps (Bushnell Core DS No Glow) to record the activity period of bird and mammal species in areas with different levels of human pressure. The study area is located in the Llanos region in Colombia. The sampling occurred from November 2020 to February of 2021 in three areas with different levels of human pressure (Fig 1). The region is highly biodiverse but also under increased pressure from agricultural expansion (Williams et al., 2020a). The areas are all immersed in a very diversified landscape with different grades of human pressure within and among the areas, ranging from isolated forest patches near human settlements and roads to continuous forest, including areas of direct influence of hydrocarbon exploitations with proximity to roads and urban settlements, as well as areas of forest, morichal and natural savannah or pastures for livestock with scattered trees and bushes.

**Figure 1.**
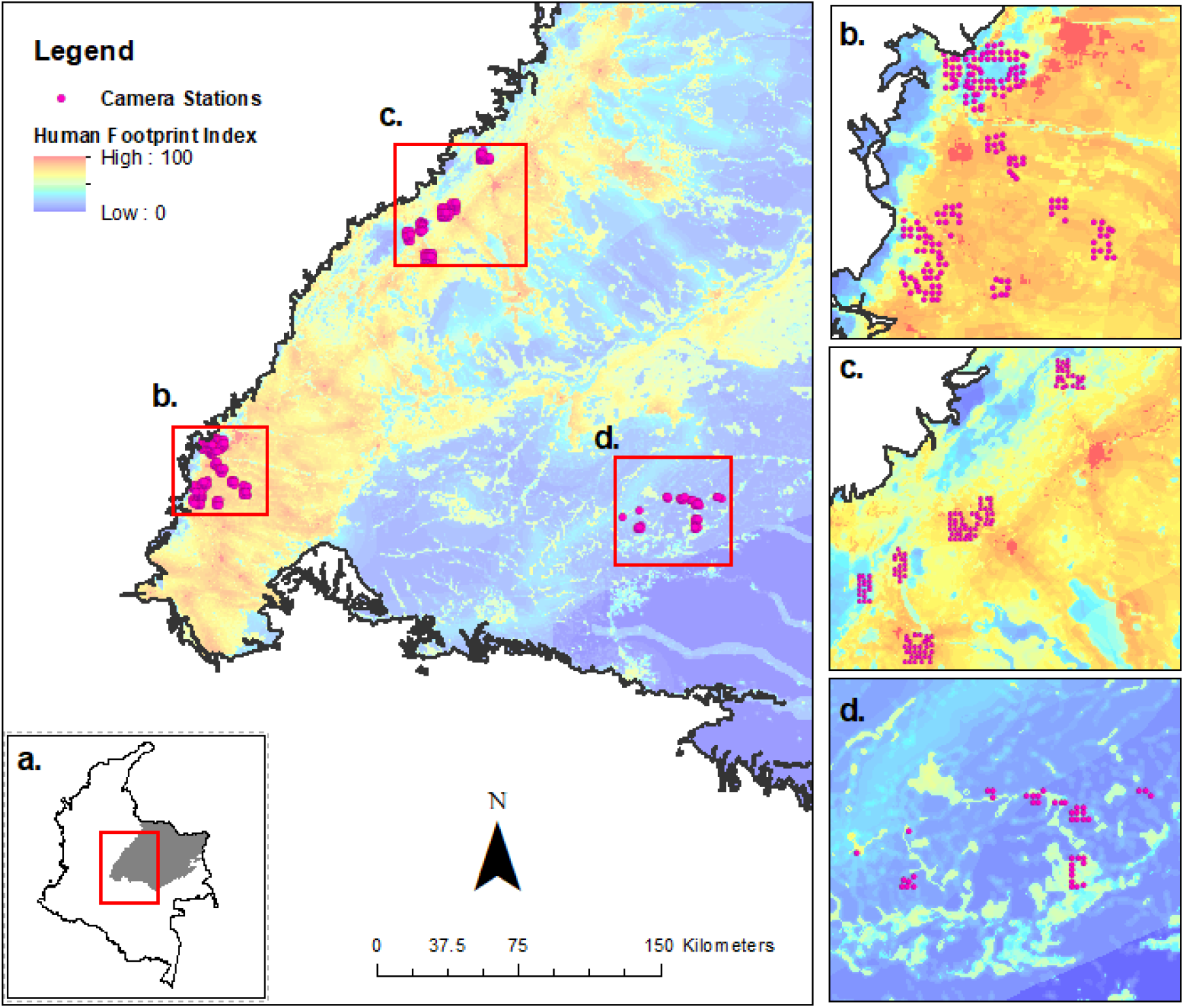
**(a)** In red the location of the study areas in Colombia. In grey the llanos region. Location of the three sites where cameras were deployed in the study area: **(b)** Meta Foothills. **(c)** Casanare Foothills. **(d)** Tillavá River. Camera trap stations are depicted by purple circles. The level of human pressure in the study area is depicted using the Legacy-adjusted Human Footprint Index (LHFI) for 2018. Red colours depict areas of high human pressure while blue colours areas of low human pressure.

We sampled each of the three sites with 367 camera stations located along a gradient of Human Footprint Index -HFI- (Correa Ayram et al., 2020); 186 in the Meta Foothills site (22-85 HFI), 138 in the Casanare Foothills site (22-91 HFI) and 43 in the Tillavá River site (8-45 HFI). Each station received one camera for a period between 16 and 96 days. Each camera trap station was placed no less than one km away from any other camera. The camera-traps were placed around 50 cm above the ground, recording photos in response to the activation of a passive infrared sensor. All images were stored and processed in the Wildlife Insights platform and specialists verified the animal identifications (Diaz-Pulido et al., 2022a, 2022b, 2022c). We used 30 minutes as a minimum time allowed between two records for the same species at the same camera, as a safety measure to avoid repeated records of the same individual. This effort resulted on a total of 16,939 camera/days and 165 species recorded; 120 birds and 45 mammals (SI Table 1).

With the data obtained from camera-trapping, we classified the records in two groups based on the time of sunset and sunrise. Records taken when the sun was up in the sky (i.e. after sunrise and before sunset) were considered “day” records, while records taken after the sunset or before sunrise were considered “night” records. Since the exact timing of sunrise slightly varies along the study area according to latitude and date, we determined each camera station's longitude, latitude, date and time and determined the exact timing of sunrise and sunset from the U.S naval observatory solar calendar (U.S. Naval Observatory, 2022). The records for each species in each sampling station were classified in day/night. Species with less than 5 captures in either the day or night were excluded from the analysis as the number of observations for these species was not enough to allow for statistical comparison. Then, using the day/night records for each one of the camera trap stations as a binomial variable (this is, every record as a Bernoulli trial), we evaluated the effect of human pressure on 11 bird and 14 mammal species, which had enough data to perform the analysis across the study area (Table 1).

**Table 1.** Main results of the regression analysis, testing the effect of human pressure on the diurnality of each mammal and bird species assessed. Significant results are in bold.

| Class | Species | Common name | Spatial Scale | Estimate | P value | Z value | Sample Size<br>(Cameras) | N. Records |
| --- | --- | --- | --- | --- | --- | --- | --- | --- |
| Mammals | *Cerdocyon thous* | Crab-eating fox | 500 | 0.03 | 0.244 | 1.164 | 56 | 293 |
|  | *Cuniculus paca* | **Paca** | 500 | -0.07 | **<<0.01** | -6.156 | 107 | 621 |
|  | *Dasyprocta fuliginosa* | **Agouti** | 1000 | -0.04 | **<<0.01** | -3.838 | 71 | 371 |
|  | *Dasybus novemcinctus* | **Nine-banded armadillo** | 500 | -0.03 | **0.046** | -1.999 | 137 | 715 |
|  | *Didelphis marsupialis* | **Common opossum** | 500 | -0.10 | **<<0.01** | -7.977 | 171 | 1030 |
|  | *Eira barbara* | **Tayra** | 500 | 0.04 | **0.006** | 2.767 | 73 | 193 |
|  | *Leopardus pardalis* | Ocelot | 1000 | 0.00 | 0.895 | 0.132 | 60 | 83 |
|  | *Mazama americana* | Red brocket | 500 | 0.02 | 0.330 | 0.974 | 41 | 117 |
|  | *Myrmecophaga tridactyla* | Giant anteater | 1000 | -0.01 | 0.324 | -0.986 | 102 | 283 |
|  | *Nasua nasua* | Ring tailed coati | 1000 | 0.04 | 0.295 | 1.048 | 19 | 29 |
|  | *Odocoileus virginianus* | White-tailed deer | 500 | -0.02 | 0.149 | -1.442 | 35 | 76 |
|  | *Pecari tajacu* | Collared peccary | 250 | 0.01 | 0.693 | 0.395 | 33 | 78 |
|  | *Philander opossum* | **Gray four-eyed opossum** | 1000 | -0.08 | **<<0.01** | -5.035 | 58 | 479 |
|  | *Tamandua tetradactyla* | **Collared anteater** | 250 | -0.03 | **<<0.01** | -3.868 | 164 | 362 |
| Birds | *Aramides cajaneus* | Grey-necked wood rail | 1000 | 0.01 | 0.115 | 1.578 | 114 | 1044 |
|  | *Arremon taciturnus* | Pectoral sparrow | 250 | -0.02 | 0.240 | -1.176 | 23 | 77 |
|  | *Catharus ustulatus* | **Swainson's thrush** | 500 | 0.10 | **<<0.01** | 4.608 | 51 | 546 |
|  | *Crypturellus cinereus* | Cinereous tinamou | 250 | -0.01 | 0.260 | -1.127 | 109 | 428 |
|  | *Crypturellus soui* | **Little tinamou** | 250 | 0.07 | **0.003** | 2.948 | 48 | 188 |
|  | *Leptotila rufaxilla* | Grey-fronted dove | 1000 | 0.02 | 0.239 | 1.176 | 124 | 1451 |
|  | *Leptotila verreauxi* | **White-tipped dove** | 1000 | 0.09 | **<<0.01** | 4.331 | 87 | 647 |
|  | *Momotus momota* | **Amazonian motmot** | 500 | 0.05 | **0.011** | 2.535 | 56 | 176 |
|  | *Myrmoborus myotherinus* | Black-faced antbird | 500 | 0.10 | 0.416 | 0.813 | 6 | 20 |
|  | *Nyctidromus albicollis* | **Pauraque** | 250 | -0.05 | **<<0.01** | -4.963 | 48 | 318 |
|  | *Vanellus chilensis* | Southern lapwing | 1000 | -0.94 | 0.317 | -1.000 | 4 | 17 |

### Human pressure proxy

We used the recently developed Colombian Legacy-adjusted Human Footprint Index (LHFI) for 2018, which is the most recent product of its kind for Colombia. It is temporarily consistent, and has a more precise spatial resolution (300 m) than any other national (Etter et al., 2011) or global human footprint map (Venter et al., 2016; Williams et al., 2020b). This index measures the level of human pressure on ecosystems and provides a baseline for monitoring human pressure on biodiversity. It was generated by the Humboldt institute (Correa Ayram et al., 2019) using the same methodology as for the one developed for the year 2015 by Correa Ayram et al., (2020). The LHFI is estimated using seven primary variables: land use type, rural population density, distance to roads, distance to settlements, fragmentation of natural vegetation, the biomass relative to natural potential, and time of intervention on ecosystems in years. All seven primary variables were estimated at 2018 and re-scaled between 0 and 5 to reflect their relative contribution to human impact and transformation. Values of 0 indicated a null contribution, while values of 5 indicated a very high contribution. Then the variables were added, and the final index was normalized between 0 and 100. See Correa Ayram et al., (2020) for a complete description of the variables and the methodology. The Human Footprint Index -HFI- (Correa Ayram et al., 2020) for each camera station was recorded.

### Data analysis

We used logistic regressions to evaluate the effect of human pressure on birds’ and mammals’ daily activity patterns, with the day/night records for each species within each camera station as a binomial response variable and the LHFI as the explanatory variable. To determine the best spatial scale for each species, four logistic models were created using the average LHFI from buffers with radius of 250, 500, 750 and 1000 m around each camera trap station, which were compared by the Akaike Information Criterion corrected for small sample sizes (AICc) (Burnham and Anderson, 2018). For each species, the model with smallest AICc value was used for the logistic regression, while the others were excluded. Since the model selection was used only to select the best spatial scale for each species, a null model was not required (Mendes et al., 2020). For the species which presented responses to human pressure, we also calculated their activity shift, here described as the difference in the model predicted diurnality between the most preserved and most disturbed areas in which the species were recorded. Finally, the activity period of all species was also plotted for visual inspection (Fig. 2), and circular metrics of diel activity, such as mean direction and Rho (i.e. a measurement of circular mean and dispersion, respectively) were calculated and reported (SI Table 2). The species’ diet, body size, activity pattern (diurnal, nocturnal or cathemeral) and whether it was a game species or not was determined based on both local expert knowledge and literature (Emmons and Feer, 1997; Hilty and Brown, 2001; IUCN, 2022; Myers et al., 2022; Ocampo et al., 2021). All analysis were performed in R version 4.2.1 (R Core Team, 2022), using the packages “lme4” (Bates et al., 2015), “circular” (Agostinelli and Lund, 2022), and ggplot2 (Wickham, 2016).

**Figure 2.**
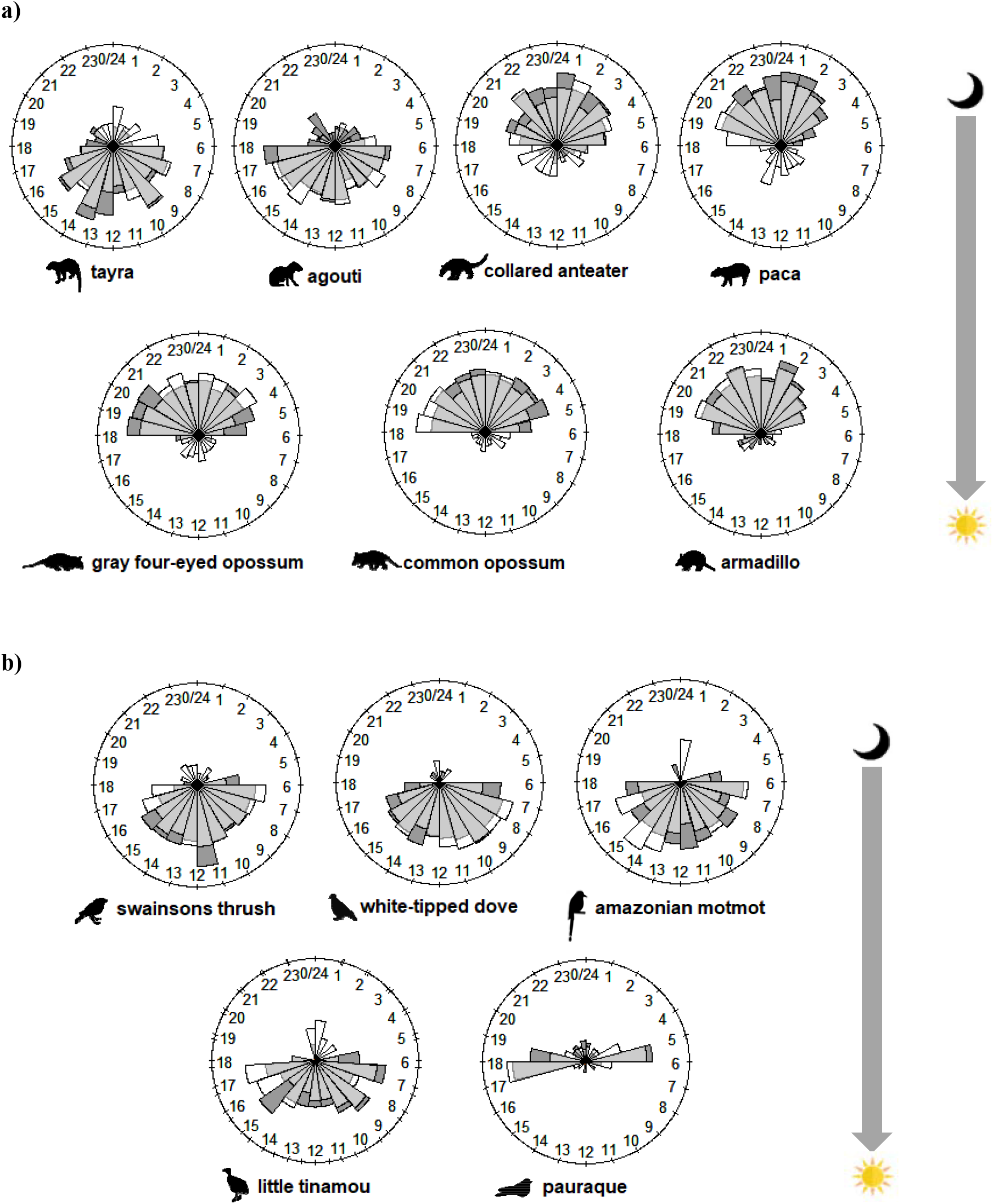
Circadian activity of **(a)** seven mammals and **(b)** five bird species in three areas in the Colombian llanos. White bars represent camaras deployed in more preserved sited, and dark grey bars represent camaras deployed in more disturbed areas. Light grey is the overlap in the circadian activity between the more preserved and more disturbed camara sites. For this figure, the location where a camara was deployed was considered disturbed when it presented a Human Footprint index above the mean, considering only the sited where camaras where deployed that had recordings of the species. The species depicted are the ones that presented statistically significant changes in diurnality based on a logistic regression. Mammal and bird silhouettes were obtained from the PhyloPic public domain database (http://phylopic.org/).

## Results

We found evidence of significant activity shifts in response to human pressure for seven out of 14 species of mammals (50%) and five of the 11 species of birds (45%) that had enough records to be assessed. For mammals, six became more nocturnal in disturbed areas while one more diurnal (Table. 1, Fig. 3a & 4a), while for birds four became more diurnal while one became more nocturnal (Table. 1, Fig. 3b & 4b). The average increase in nocturnality for mammals was 9.5% while the average increase in diurnality for birds was 7%. However, we must clarify that by “becoming more nocturnal” or “becoming more diurnal”, we did not mean that a species shifted its peak of activity from a period to another. Instead, it means that although the species main activity is still occurring during its natural activity period (i.e. night for nocturnal species and day for diurnal species), the proportions of the day/night records have changed (Mendes et al., 2020).

**Figure 3.**
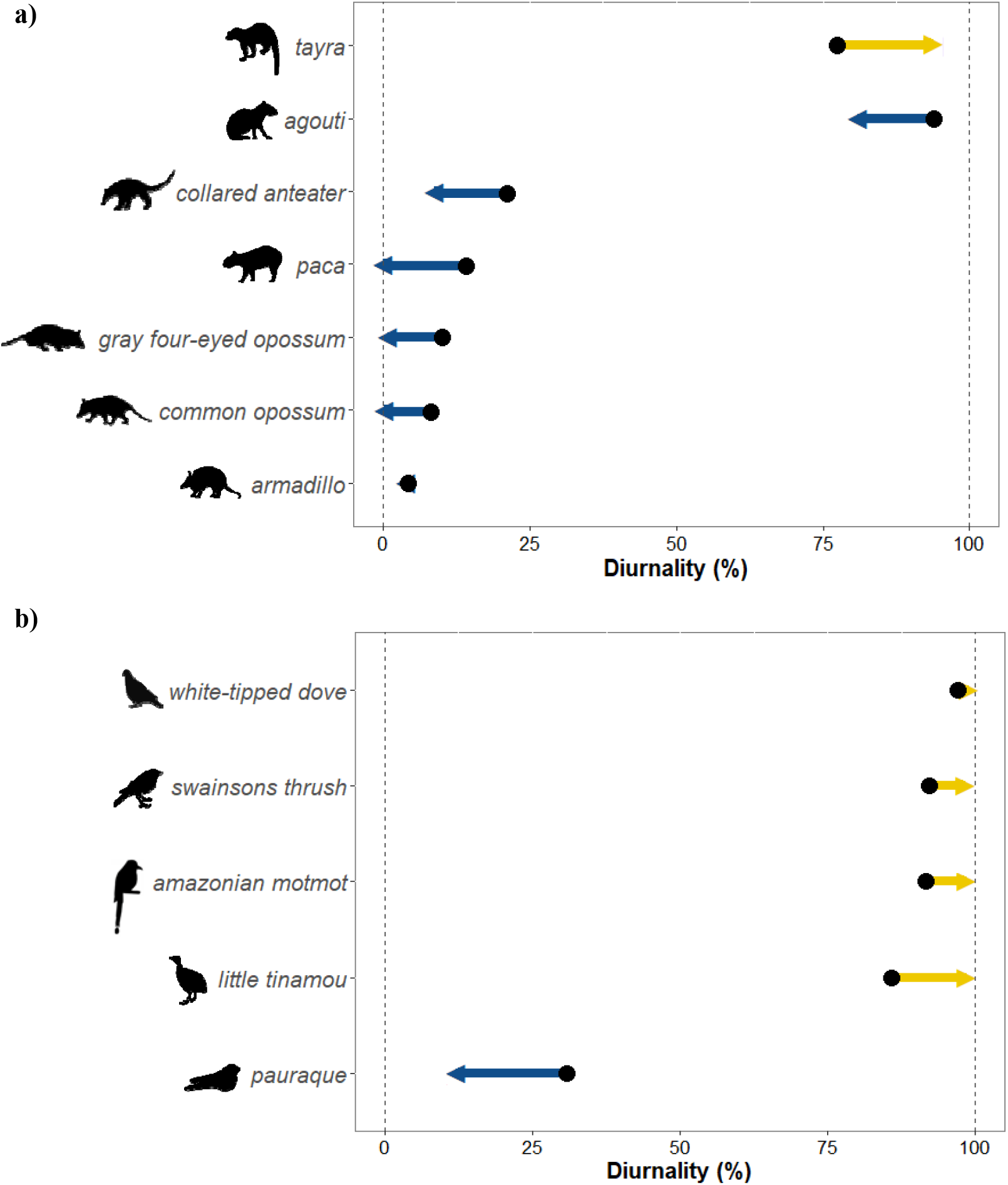
Shifts in diurnality of **(a)** seven mammals and **(b)** five bird species in response to human pressure in three sites in the Colombian llanos, from the most preserved sites (black point) to the most disturbed sites (arrow point) in which each species was recorded. Arrows to the right represent shifts toward diurnality, while arrows to the left represent shifts toward nocturnality. The species depicted are the ones that presented statistically significant changes in diurnality based on a logistic regression. Mammal and bird silhouettes were obtained from the PhyloPic public domain database (http://phylopic.org/).

**Figure 4.**
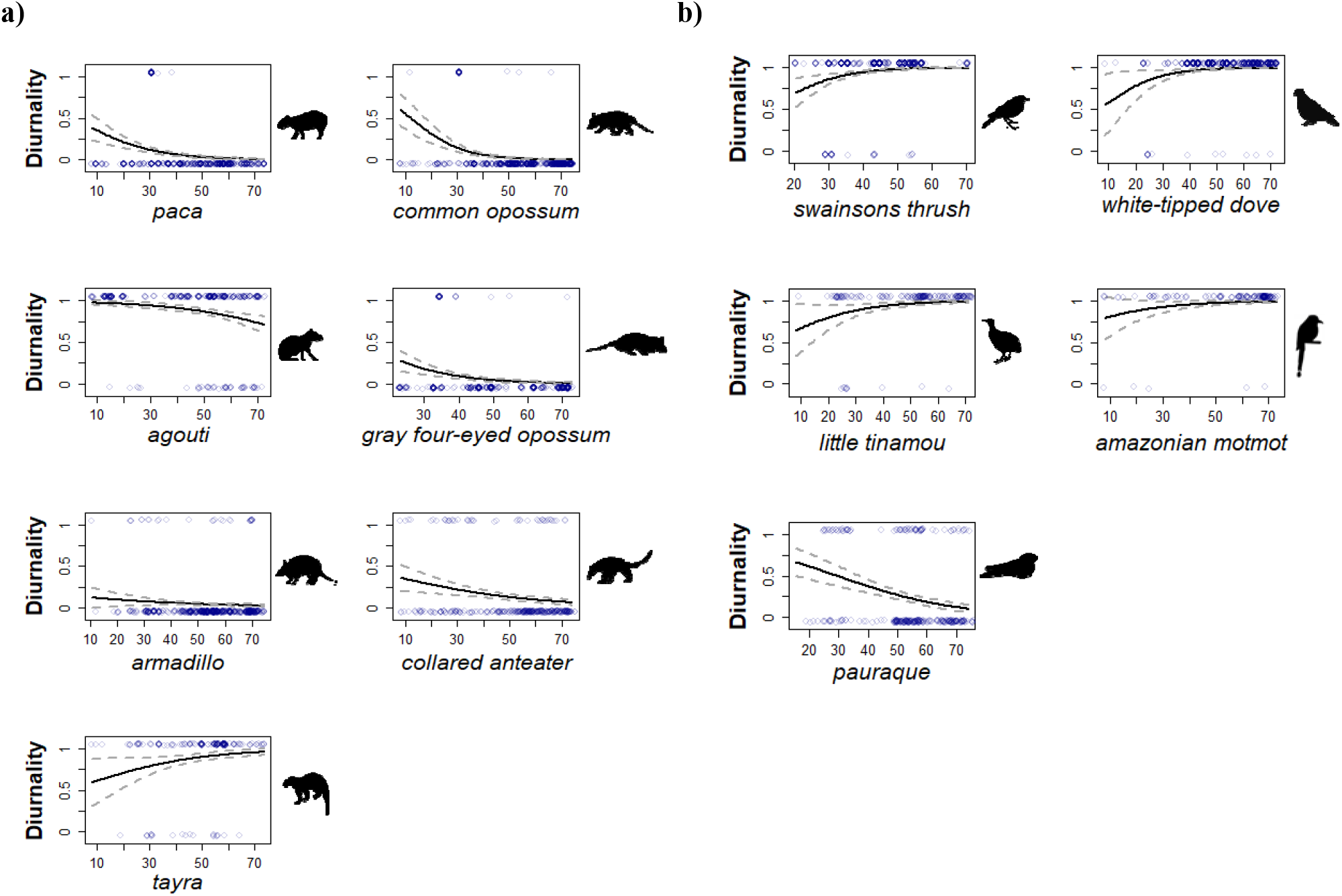
Changes in diurnal activity for **(a)** seven mammals and **(b)** five bird species with increases in human pressure. The human pressure (x-axis) was measured using the Legacy-adjusted Human Footprint Index (LHFI) for 2018 of each site as a proxy (Correa Ayram et al., 2019). The species depicted are the ones that presented statistically significant changes in diurnality based on a logistic regression. Mammal and bird silhouettes were obtained from the PhyloPic public domain database (http://phylopic.org/).

The bird species which responded to human pressure by becoming more diurnal were the swainson’s thrush (*Catharus ustulatus*), the little tinamou (*Crypturellus soui*), the white-tipped dove (*Leptotila verreauxi*) and the amazonian motmot (*Momotus momota*). The pauraque (*Nyctidromus albicollis*) was the only bird species that responded to human pressure by becoming more nocturnal. Other six bird species did not present detectable activity shifts in response to human pressure. The grey-necked wood rail (*Aramides cajaneus*), the pectoral sparrow (*Arremon taciturnus*), the cinereous tinamou (*Crypturellus cinereus*), the grey-fronted dove (*Leptotila rufaxilla*), the black-faced antbird (*Myrmoborus myotherinus*) and the southern lapwing (*Vanellus chilensis*).

In relation to mammals, the species which responded to human pressure by becoming more nocturnal were the paca (*Cuniculus paca*), the agouti (*Dasyprocta fuliginosa*), the nine-banded armadillo (*Dasypus novemcinctus*), the common opossum (*Didelphis marsupialis*), the grey four-eyed opossum (*Philander opossum*) and the collared anteater (*Tamandua tetradactyla*). Three of these species (agouti, paca and armadillo) are heavily hunted or persecuted. The tayra (*Eira barbara*) was the only mammal species that responded to human pressure by becoming more diurnal. Other seven species, the crab-eating fox (*Cerdocyon thous*), the ocelot (*Leopardus pardalis*), the red brocket (*Mazama americana*), the giant anteater (*Myrmecophaga tridactyla*), the ring-tailed coati (*Nasua nasua*), the white-tailed deer (*Odocoileus virginianus*) and the collared peccary (*Dicotyles tajacu*), did not present detectable activity shifts in response to human pressure.

## Discussion

We found evidence of significant activity shifts in response to human pressure for 45% of the bird species assessed, with four of five shifts towards more diurnal activity patterns. The four species exhibiting diurnal shifts were already diurnal and one is a game species (little tinamou; *Crypturellus soui*) and the pauraque (*Nyctidromus albicollis*) shifted from cathemeral in more intact areas towards nocturnality. The increases in diurnality may be a response to increased foraging required to counter the decreases in food availability in degraded habitats [e.g. due to logging, edges, fragmentation (Fontúrbel et al., 2021; Redpath, 1995; Saunders, 1980; Smetzer et al., 2022)]. Degraded habitats may also influence vertebrates activity patterns because they have higher variation in microclimate, which can affect other environmental factors such as sunlight (Bennie et al., 2014; Ewers and Banks-Leite, 2013; Fontúrbel et al., 2021; Schmidt et al., 2017), insect prey (Checa et al., 2014; Outhwaite et al., 2022) and other key resources like flowering, nectar production and fruit ripening (Jackson, 1966; Vitasse et al., 2021).

Three of the six bird species that didn’t show a statistically significant shift in daily activity patterns (black-faced antbird, southern lapwing and pectoral sparrow) have few captures. The grey-necked wood rail and the grey-fronted dove had non-significant shifts toward diurnality with increased human pressure. The cinereous tinamou had a non-significant increase in nocturnality with human influence, opposite to the significant shift found in the little tinamou. These two tinamous are sympatric, morphologically similar, and closely related ground-dwelling birds, which may generate a pressure for temporal niche partitioning (Dias et al., 2016; Negret et al., 2015b) that may be reinforced in degraded habitats with lower food availability.

In relation to mammals, we found that 50% of the species assessed showed activity shifts in response to human pressure. These results align with the findings of other studies that show changes in activity patterns for mammals species with increased human pressure (Bennie et al., 2014; Gaynor et al., 2018; Mendes et al., 2020). From the seven mammal species which had a significant activity shift, four are nocturnal and three of those are also game species. All of them had shifts towards a more nocturnal behaviour. From the three diurnal species which presented activity pattern shifts, the agouti and the Common opossum, which are hunted, became more nocturnal when in disturbed areas, while the tayra became more diurnal in areas with higher human pressure. The Tayra is a generalist species that is able to subsist on second growth forest and agricultural mosaics without forest cover, and may subsidize their diet taking advantage of domesticated animals such as chicken (Meneguetti et al., 2014; Michalski and Peres, 2005). These characteristics could explain its shift towards diurnality in areas where there are anthropogenic sources of food that can be opportunistically exploited during daytime.

Four of the seven mammal species that didn’t show a statistically significant shift in daily activity patterns (ocelot, ring tailed coati, white-tailed deer and collared peccary), had a low number of records which could have influenced the explanatory power of those models. The giant anteater, a species that is occasionally killed or captured and that is classified as Vulnerable (Bertassoni and Abba, 2014; Emmons and Feer, 1997) had an increase in nocturnality in areas with higher human pressure as was the case for the nocturnal mammal species that had significant changes in daily activity patterns. The crab-eating fox and red brocket deer had an increase in diurnality in areas with higher human pressure. These two species can be found in degraded landscapes and have generalist diets (Emmons and Feer, 1997), which suggests that they might be less affected by human pressure than other species.

The effect of human pressure on vertebrate species goes beyond impacting species richness and abundance. Our results show that increased human pressure can have a marked impact in the activity pattern of multiple vertebrate groups. Particularly, we found that 45% of the bird species and 50% of the mammal species assessed had shifts in their activity pattern with increases in human pressure. Bird species had a general shift towards diurnality, while mammal species towards nocturnality, revealing that different vertebrate groups can react distinctly to increased human presence and activity. There is a need for more research on the impact of human pressure on vertebrates’ behaviour, as such changes could result in marked shifts away from natural patterns, with consequences for fitness, population persistence, community interactions, and evolution (Chace and Walsh, 2006; Gaynor et al., 2018; Lin et al., 2012). The availability of camera trap information is increasing exponentially, but usually the only information reported is in relation to species’ community composition and population densities (Antunes et al., 2022; Negret et al., 2015a). Our analysis highlights the fact that many behavioural characteristics, such as species’ daily activity, may vary between species, individuals and sites and that this variation can have impacts on the ecology and conservation of these species. Future studies should focus on the impact of human pressure on the behavioural patterns of vertebrates (Gaynor et al., 2018; Sih et al., 2011), especially for birds and other groups for which information in this regard is scarce (Bonier et al., 2007; Chace and Walsh, 2006; Fontúrbel et al., 2021; Smetzer et al., 2022) and for regions that are underrepresented. Our findings are a step in this direction.

## Acknowledgments

We would like to acknowledge the people from the Maron Ecology & C onservation Policy L ab and the Ecological C ascades L ab for helpful discussion on the interpretation of the results and to C. Rojano, L. Miranda y R. Á vila (Fundación C unaguaro), A. L opera, F. Hernández, R. Fernández y R. Rodríguez (C orporación Gaica), Á. E. C áceres-Gómez, N. Ferro-Muñoz. F. C áceres-Gómez (B iotica C onsultores) for the field work, and S. Pérez for the revision of the bird species lists. We thank Ecopetrol S.A. and Instituto de Investigación de Recursos Biológicos Alexander Von Humboldt for financing the study within the framework of the FIBRAS Agreement.

## Supplementary material

**SI Table 1.**
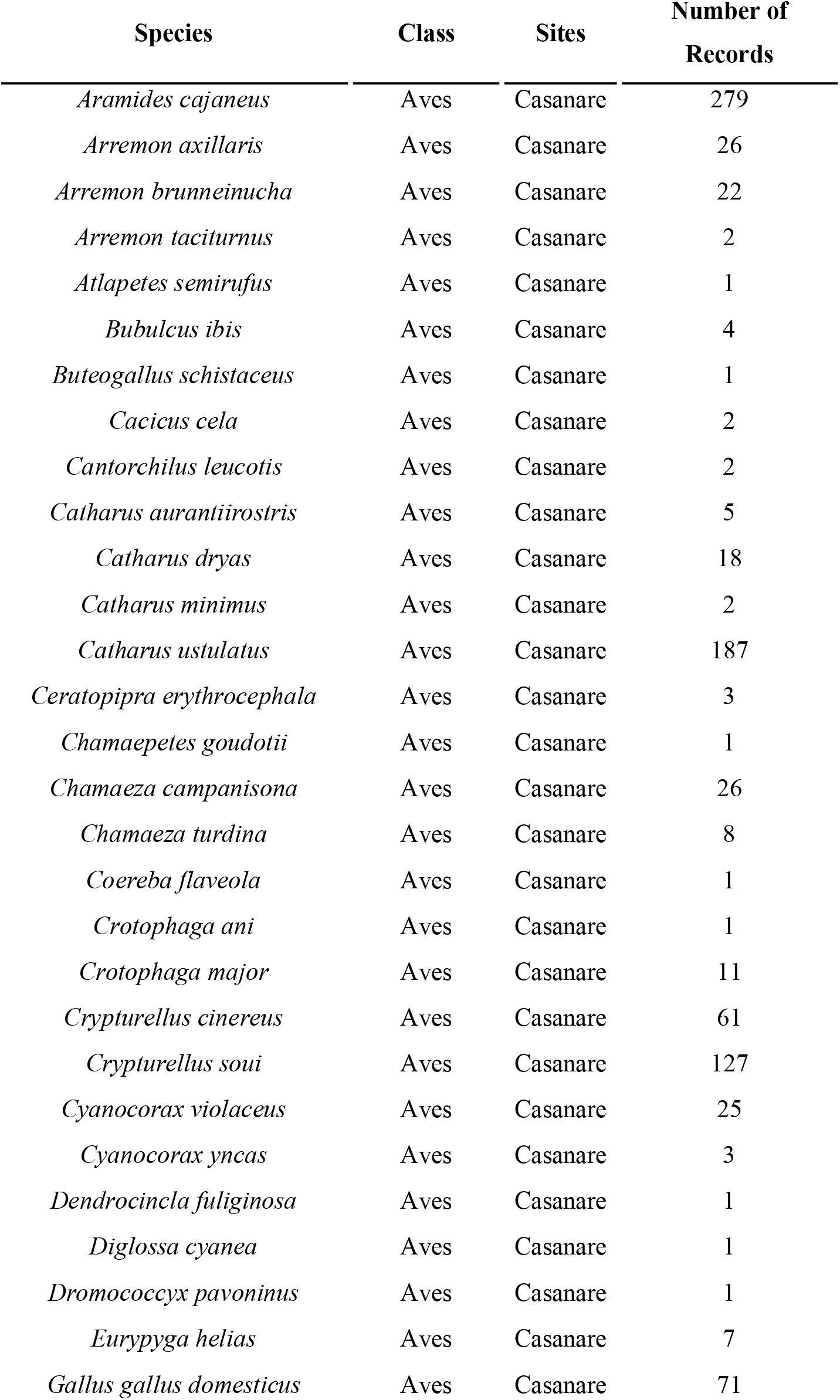
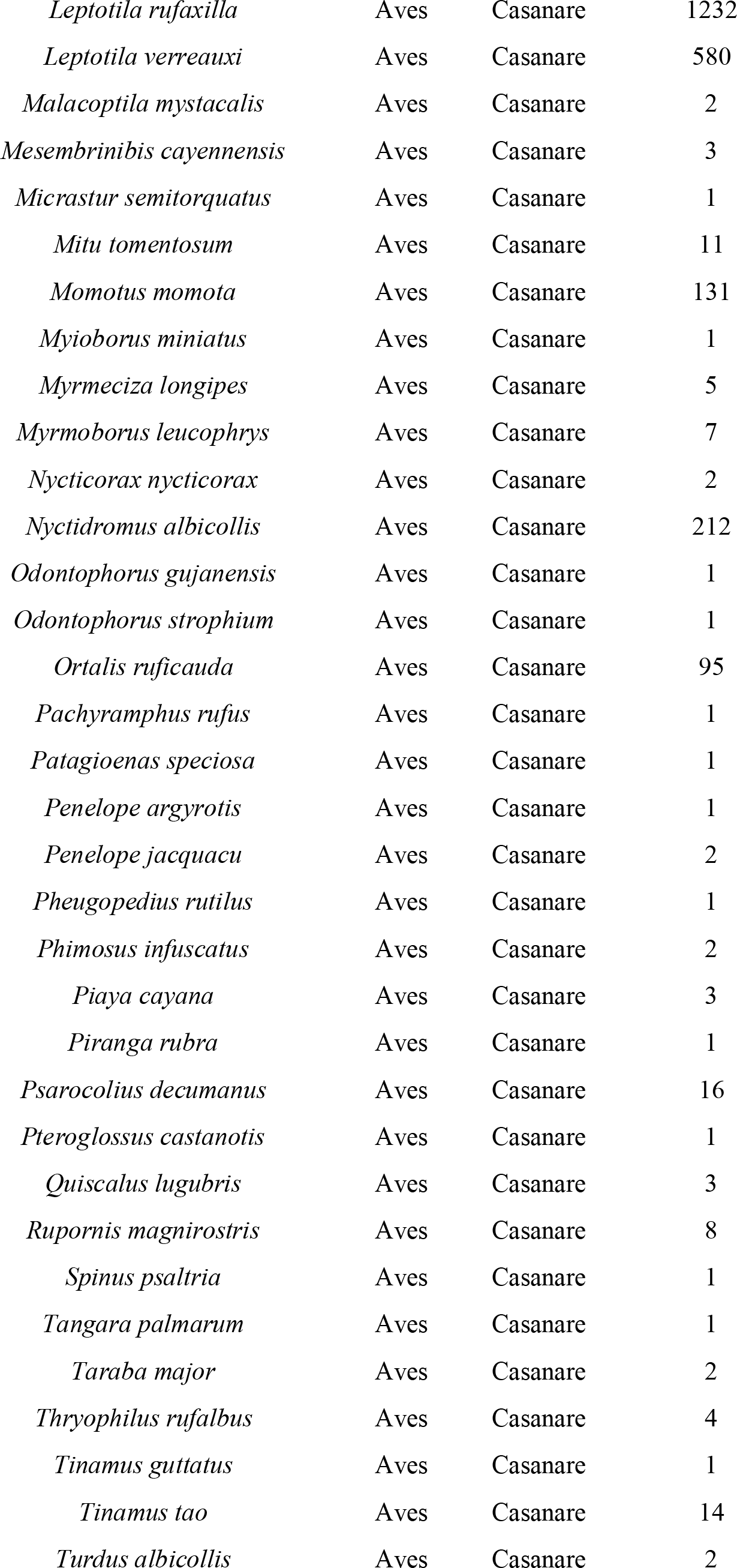
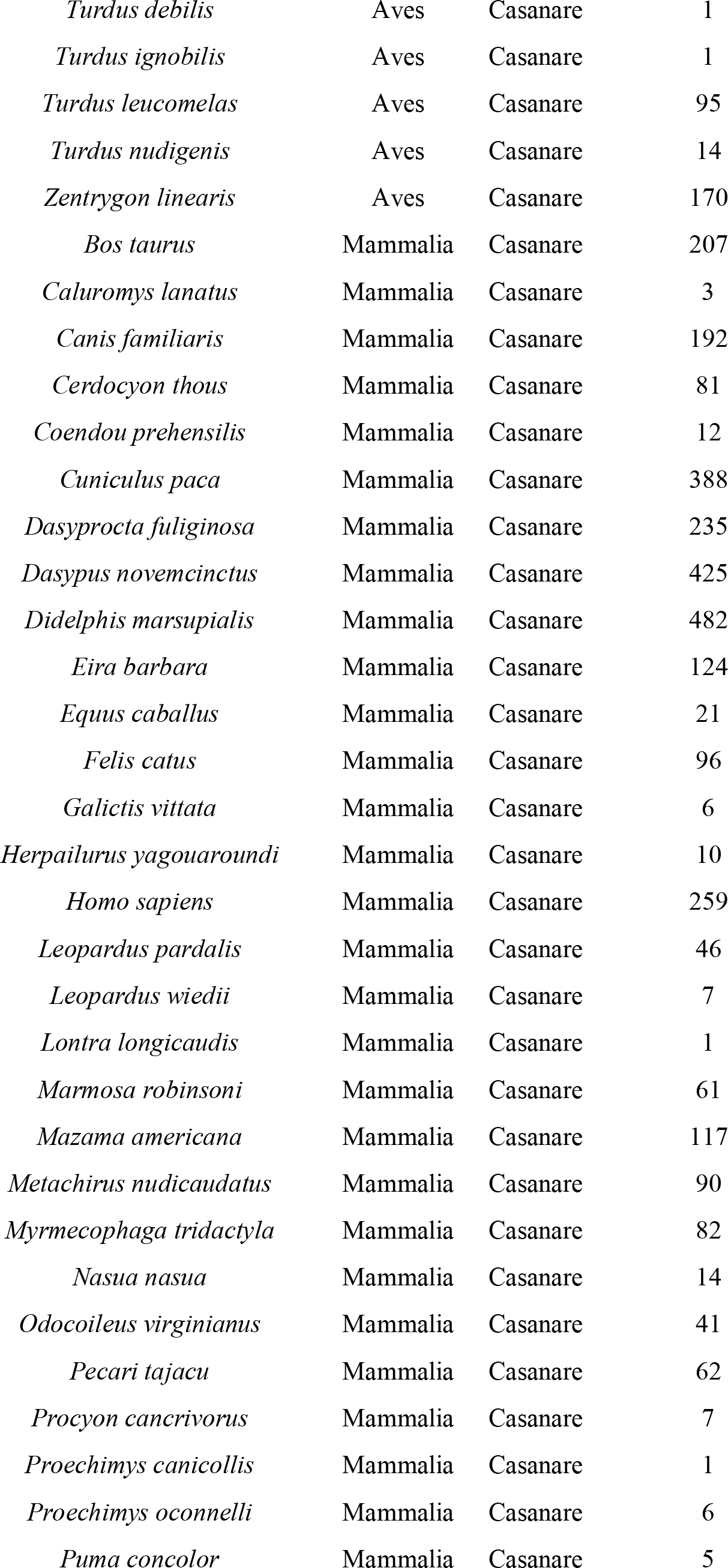
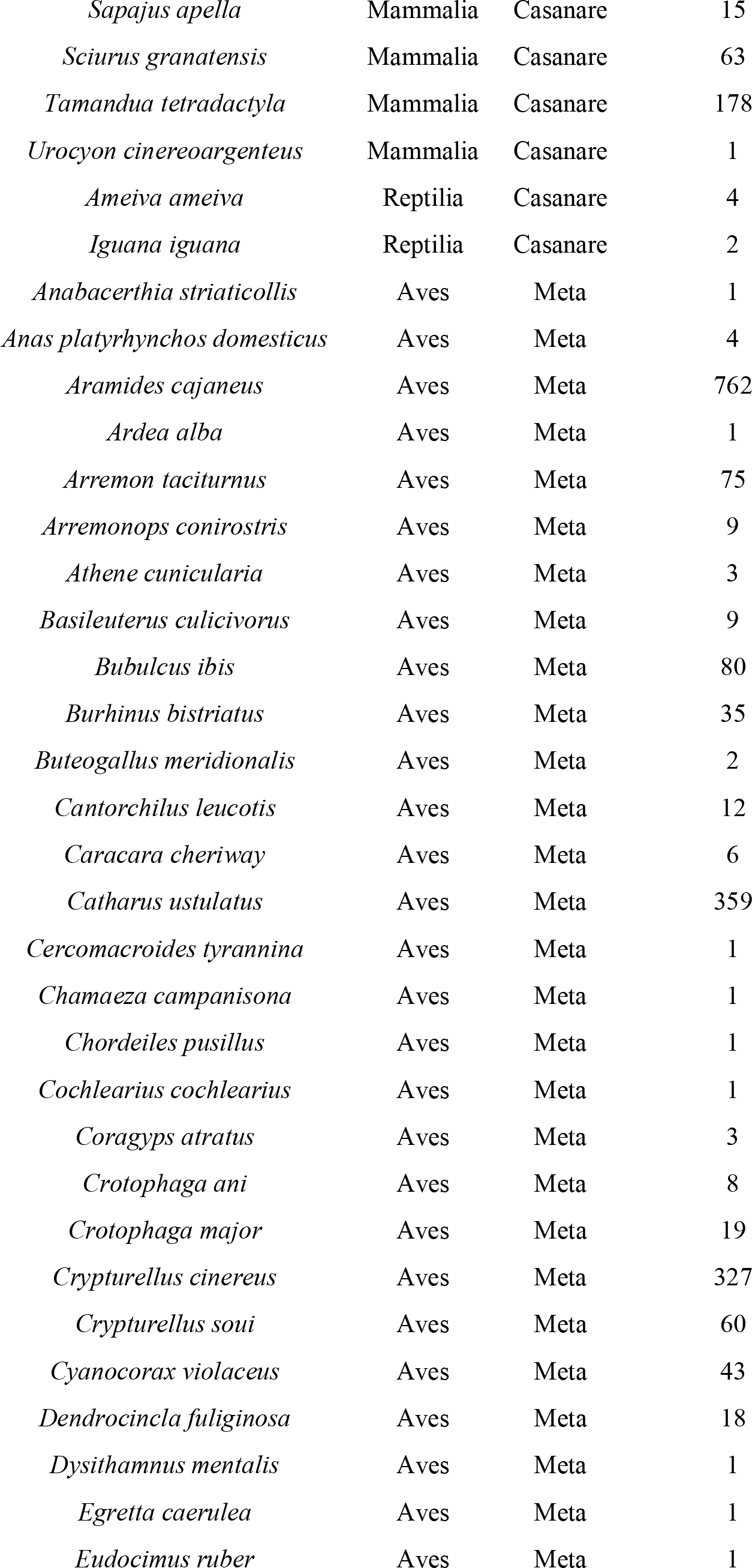
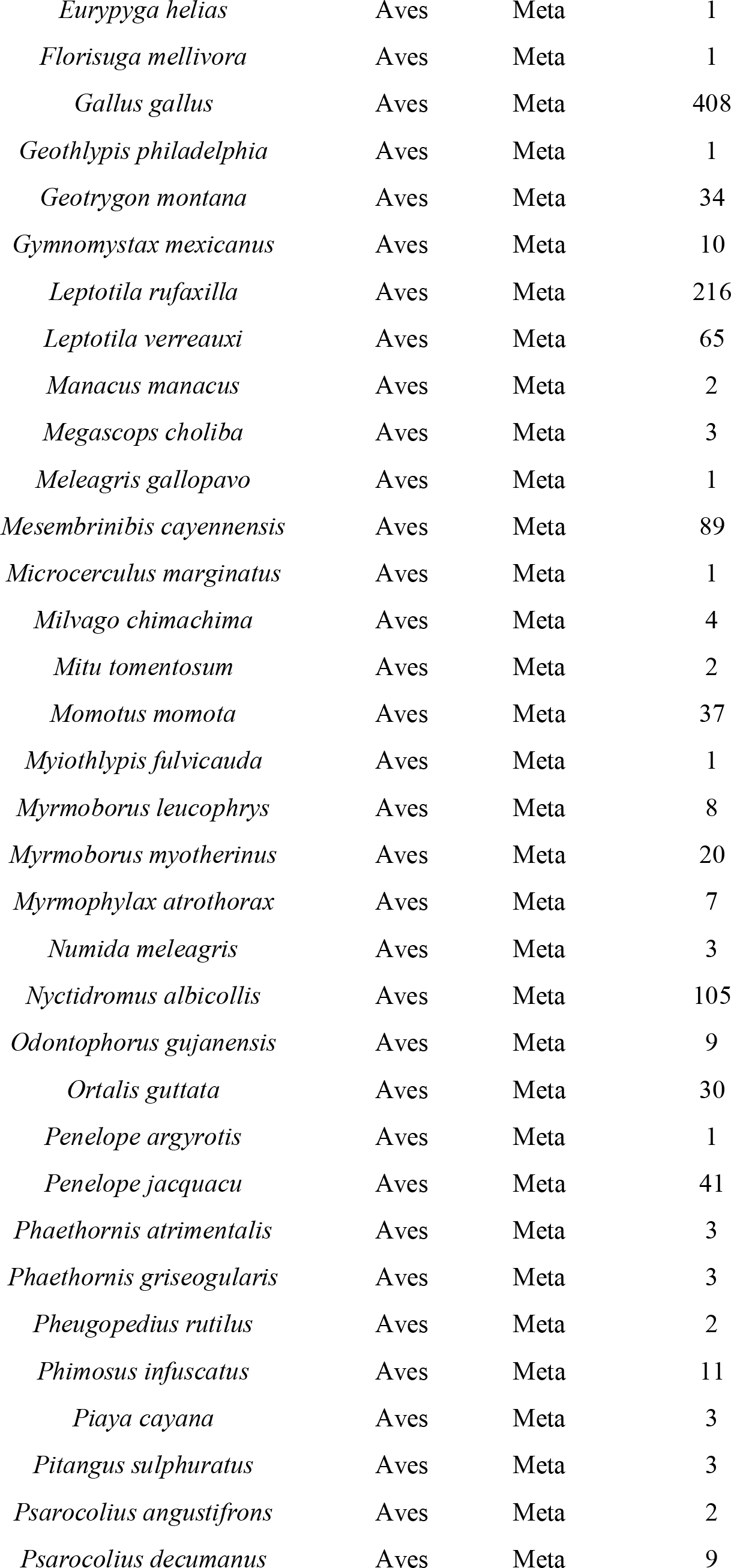
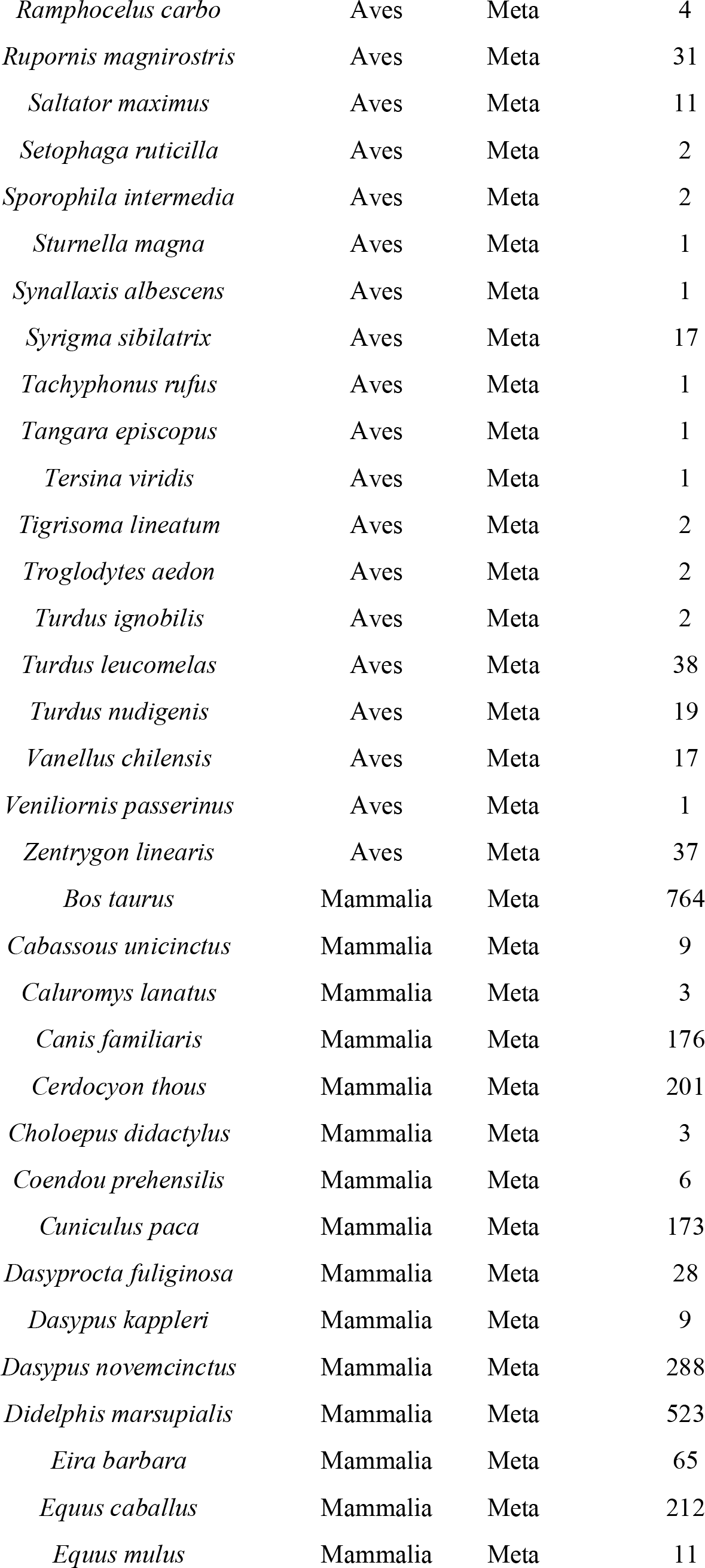
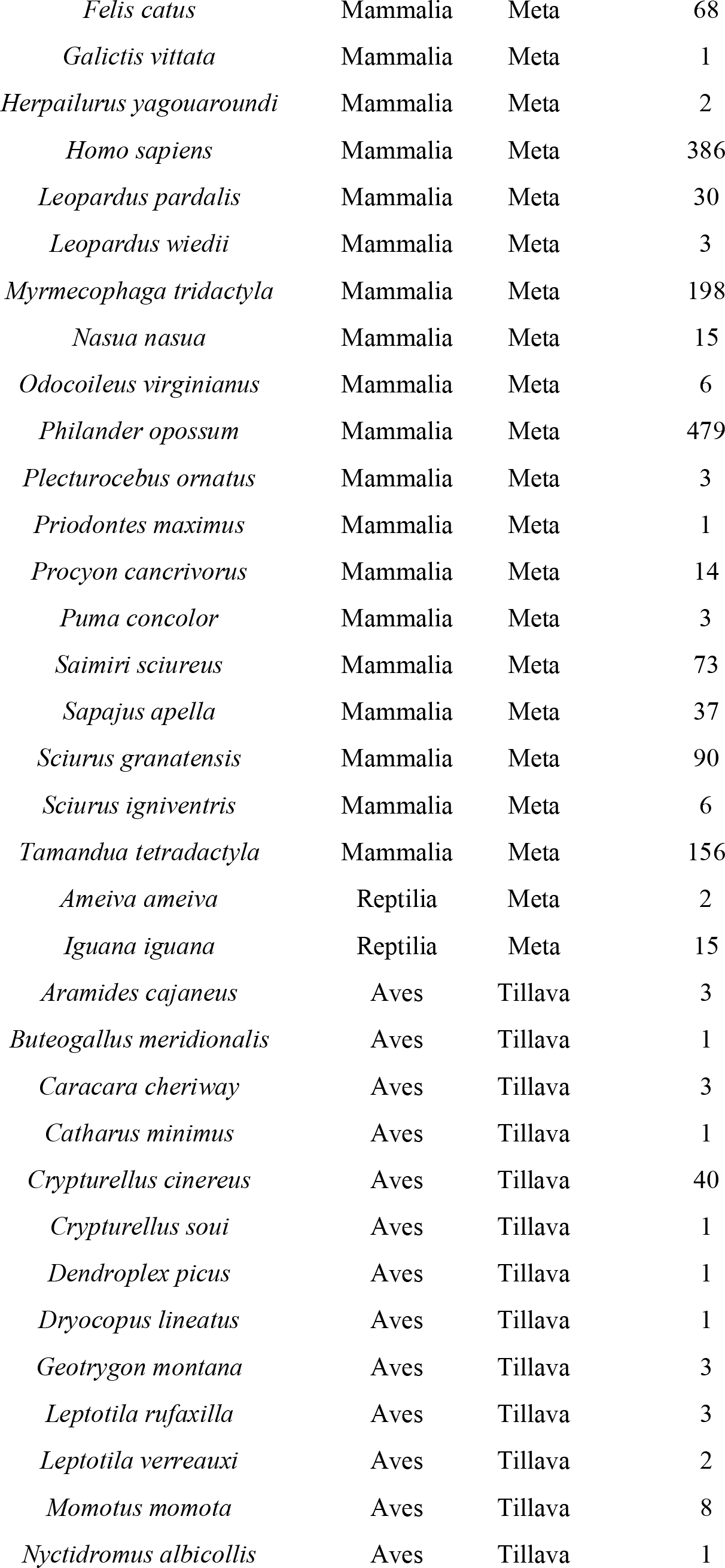
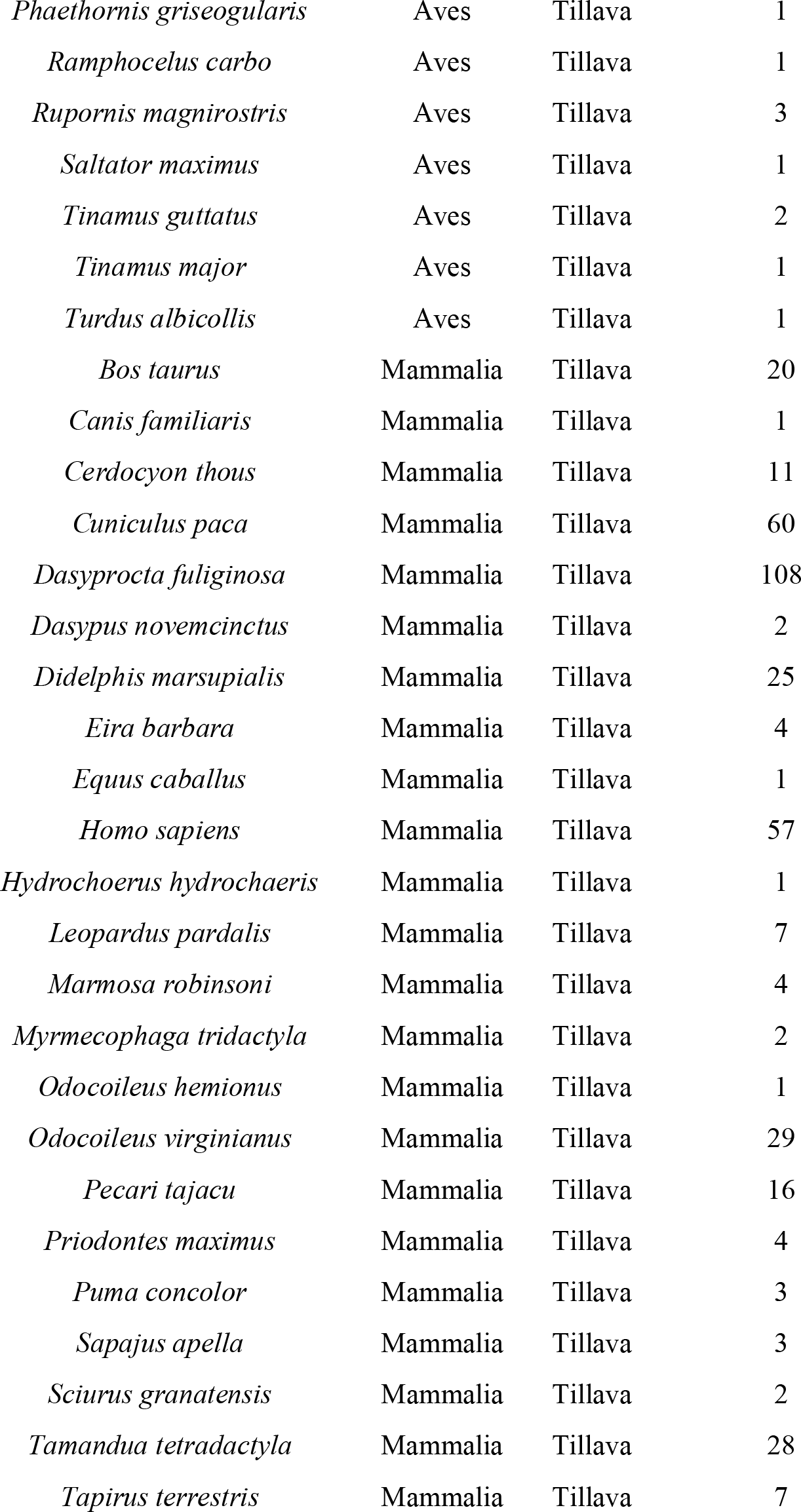
Mammal, bird and reptile species recorded with the camera traps in each of the three sites (Casanare, Meta and Tillava) in the Llanos region in Colombia.

**SI Table 2.**
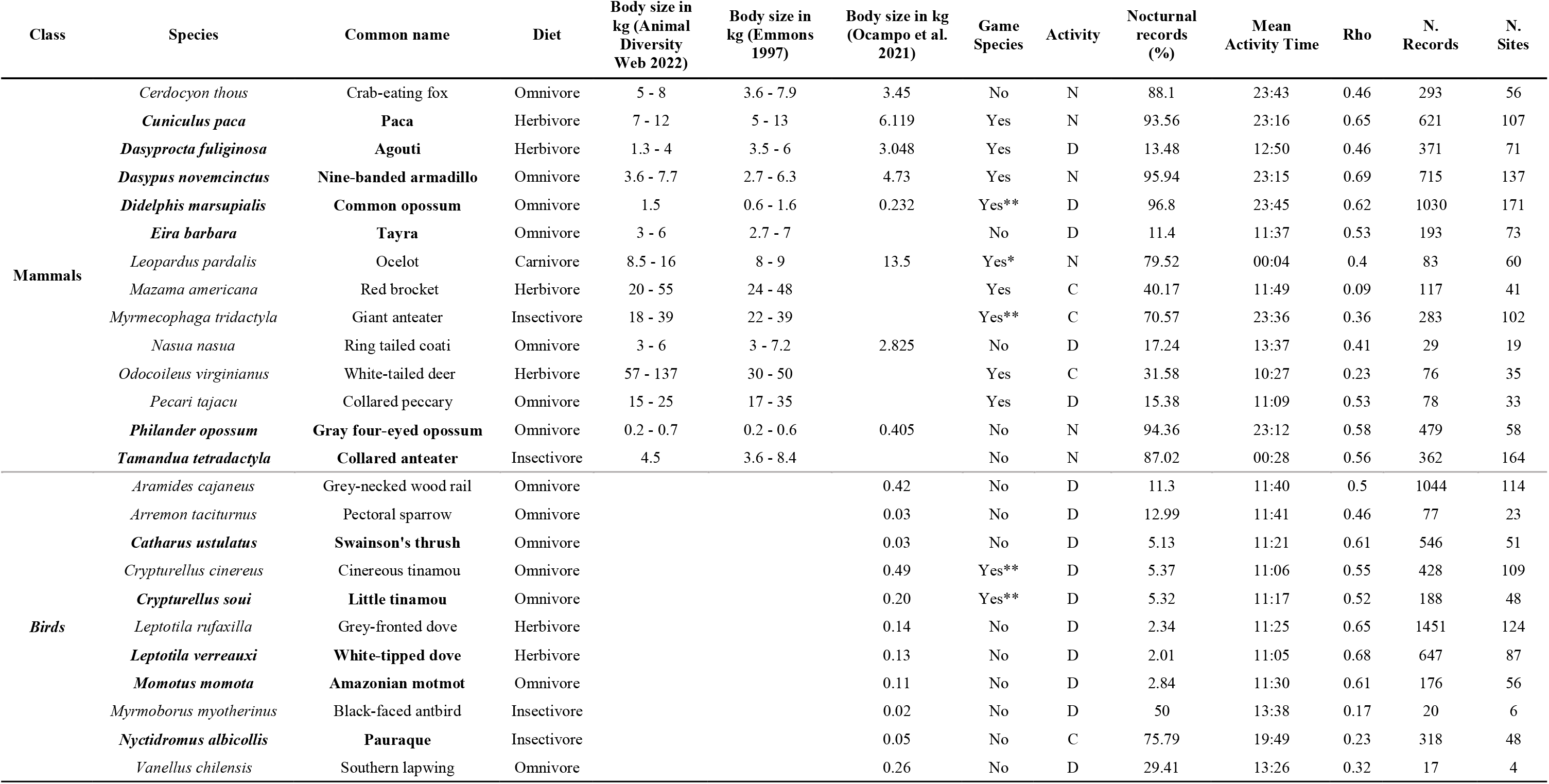
Diel activity patterns, diet, body size and hunting pressure of the 15 mammals and 11 bird species. The mean activity time is a circular statistics version of the arithmetic mean, while the Rho is a circular statistics descriptor of dispersion. Activity is abbreviated as “D” for diurnal species, “N” for nocturnal and “C” for Cathemeral. The Ocelots are hunted not for their meat but for their fur or as retaliation due to killing of domestic animals “*”. Tinamous, the giant ant eater and the common opossum can be opportunistically hunted but are not heavily persecuted “**”.

